# Ascites in malignant ovarian cancer confer ovarian cancer cells chemoresistance to platinum. Combination treatment with crizotinib and 2-hydroxyestradiol restores platinum sensitivity

**DOI:** 10.1101/2023.09.24.559160

**Authors:** Yifat Koren Carmi, Abed Agbarya, Hazem Khamaisi, Raymond Farah, Yelena Shechtman, Roman Korobochka, Jacob Gopas, Jamal Mahajna

**Affiliations:** Department of Nutrition and Natural Products, Migal – Galilee Research Institute, Kiryat Shmona, Israel; Shraga Segal Department of Microbiology, Immunology and Genetics, Ben Gurion University of the Negev, and Department of Oncology, Soroka University Medical Center, Beer Sheva, Israel; Oncology Department, Bnai Zion MC, Haifa, Israel; Department of Internal Medicine, Ziv Medical Center, Safed, Israel; Department of Biotechnology, Tel-Hai College, Kiryat Shmona, Israel

**Keywords:** Ovarian cancer, platinum chemoresistance, tumor microenvironment, 2-hydroxyestradiol, Crizotinib

## Abstract

Ovarian cancer (OC), the second most common form of gynecologic malignancy, has a poor prognosis and is often discovered in the late stages. Platinum-based chemotherapy is the first line of therapy. Nevertheless, treatment OC has proven challenging due to toxicity and the development of acquired resistance to therapy. Tumor microenvironment (TME) has been associated with platinum chemoresistance.

Malignant ascites has been used as OC tumor microenvironment and its ability to induce platinum chemoresistance has been investigated.

Our results suggest that exposure to OC ascites induces platinum chemoresistance in 11 of 13 cases (85%) on OC cells. In contrast, 75% of cirrhotic ascites (3 of 4) failed to confer platinum chemoresistance to OC cells. Cytokine array analysis revealed that IL -6 and to a lesser extent HGF were enriched in OC ascites, whereas IL -22 was enriched in cirrhotic ascites. Pharmaceutical inhibitors targeting the IL -6/ JAK pathway were mildly effective in overcoming platinum chemoresistance induced by malignant ascites. In contrast, crizotinib, an HGF/c-MET inhibitor, and 2-hydroxyestradiol (2HE2) were effective in restoring platinum chemosensitivity to OC.

Our results demonstrate the importance of OC ascites in supporting platinum chemoresistance and the potential of combination therapy to restore chemosensitivity of OC cells.

## 1 Introduction

Epithelial ovarian cancer (EOC), makes up 90% of ovarian cancer (OC) cases. There are five different types of EOC: mucinous, low and high-grade serous ovarian cancer (HGSOC), endometroid, clear cells, and unspecified. HGSOC is the most prevalent and deadly type (1). OC is the second most common form of gynecological malignancy and has a poor 5 years survival rate (2). Because of late-stage diagnosis, HGSOC accounts for the majority of OC death cases (1, 3). The standard of care for OC is platinum-based chemotherapy. The management of patients with OC has, however, been complicated by toxicity and acquired resistance (4, 5). Chemoresistance frequently occurs in OC patients due to a number of mechanisms, including reduced platinum accumulation within the cancer cells, increased drug inactivation by metallothionein and glutathione, and increased DNA-repair activity (6, 7). A number of studies have linked the tumor microenvironment (TME) to cancer chemoresistance in addition to chemoresistance caused by alterations within cancer cells (6–9) .

TME can be separated into cellular and soluble parts. Multiple cell types, including immune cells, fibroblasts, adipocytes, and mesenchymal stem cells (MSC), constitute the cellular compartment of TME. The majority of the substances found in the soluble compartment are factors secreted by the TME’s cellular compartment.

The peritoneum is where OC cells likely metastasize and form implants on the omentum. Ascites in OC are produced via the increased secretion of vascular endothelial growth factor (VEGF) by the cancer cells, which makes the blood vessels more permeable and/or obstruction of lymph drainage by the tumor. Cancerous cell spheroids, hematopoietic cells, adipocytes, fibroblasts, mesothelial cells, and MSC produced from bone marrow are some of the cellular components of ascites (10, 11). Non-cellular constituents, such as extracellular matrix (ECM) components, extracellular vesicles, exosomes, cytokines, growth factors, metabolites, and lipid synthesis mediators have been identified in ascites (11, 12).

Malignant ascites is a common presentation in more than one-third of OC patients. The microenvironment in OC malignant ascites is pro-inflammatory and tumor-promoting, which has been linked to metastasis and chemoresistance (3) (13–17). Malignant ascites serves as a reservoir for a diverse mixture of soluble factors, metabolites, and cellular components. Development of ascites is a major indication of disease recurrence (18, 19). Ascites formation onset and progression is linked to patients’ declining quality of life and a poor prognosis (18). However, it is not clear whether ascites is an indication of chemotherapy failure or if ascites per se is associated with the emergence of chemoresistance. Tumor cells within the ascites were shown to exhibit cancer stem-like characteristics that possess increased resistance to treatments and the capacity for distal metastatic spread in recurrent disease (20). Ascites microenvironment is a significant cause of morbidity and mortality in OC patients.

Ascites has the potential to be a predictor of drug response, as a substrate for assessing drug efficacy, and possibly as a tool to identify early indications of chemoresistance, disease progression, and therapy modifications. Therefore, we analyzed the ascites unique tumor microenvironment to investigate platinum chemoresistance and potential measures to overcome it.

As previously reported, cellular and soluble TME compartments play a role in conferring platinum chemoresistance to OC cells (21) (22) (23). Those findings showed that direct co-culture of OC with MSC resulted in conferring chemoresistance to therapeutic drugs such as paclitaxel, colchicine, and platinum compounds, alongside blocking of ERK1/2 activation (21). It was demonstrated that the inclusion of fisetin and other flavonoids to a platinum drug, restored platinum drug sensitivity to OC cells co-cultured with MSC, accompanied by reactivation of ERK1/2 (21). Furthermore, 2-hydroxyestardiol (2HE2), an estradiol metabolite that serves as a prodrug converting to 2-methoxyestradil (2ME2), was effective in restoring platinum sensitivity to OC cells growing in direct interaction with MSC (24). Moreover, we reported that conditioned media (CM) derived from human stromal cells (HS-5) and tumor-activated HS-5 (TA-HS-5) were effective in conferring platinum chemoresistance to OC cells (23). The soluble factors IL-6, and hepatocyte growth factor (HGF) were implicated in mediating platinum chemoresistance that could be partially or completely overcome by pharmaceutical inhibitors targeting the JAK/STAT and c-MET signaling pathway, respectively (23).

The aim of our study is to investigate the role and clinical significance of TME in chemoresistance of OC to monitor the ability of liquid ascites fluids to affect platinum sensitivity of OC cells and to explore means to restore platinum sensitivity to OC cells.

In this study, we examined the ability of ascites fluids derived from OC patients in comparison to non-malignant cirrhosis ascites to affect platinum sensitivity of OC cells. Contrary to the majority of cirrhosis ascites, the majority of OC ascites confer platinum chemoresistance to OC cells. The cytokine IL-6, as well as HGF and to a lesser degree G-CSF, were enriched in OC ascites, whereas IL-22 was enriched in cirrhosis ascites. Pharmaceutical inhibitors that target the JAK signaling pathway have a moderate activity in overcoming platinum chemoresistance mediated by malignant ascites. Additionally, 2HE2 and Crizotinib, c-MET inhibitors, were active in restoring platinum chemosensitivity to OC. Our findings demonstrated the importance of OC ascites in promoting platinum chemoresistance as well as the potential of combination therapy including JAK and c-MET inhibitors and 2HE2 in restoring platinum chemosensitivity to OC cells.

## 2 Materials and Methods

### 2.1 Chemicals and reagents

Most chemicals were obtained from Sigma Aldrich Israel Ltd. (Rehovot, Israel), otherwise the vendor is specified. 2HE2 was purchased from Cayman Chemical (Ann Arbor, Michigan, USA). Ruxolitinib, Tofacitinib, Dasatinib and Crizotinib obtained from Selleck Chemicals LLC (Houston, TX, USA) and platinum IV prodrug RJY13 (25).

### 2.2 Cell lines

Human cisplatin-resistant and RJY-13-sensitive cells OC A2780cisR (obtained from the American Type Culture Collection, VA, USA), were cultured in RPMI 1640 complete medium supplemented with 10% (w/v) fetal bovine serum (Biological Industries, Israel), 1% (w/v) L-glutamine, 100 units/ml penicillin, and 0.1 mg/ml streptomycin. The human and murine MSC lines, HS-5 and MS-5, respectively were maintained under the same conditions. All cell lines were grown at 37°C in a humidified atmosphere with 5% CO_2_.

### 2.3 Trypan blue exclusion assay

Cells (2 x 10^5^/well) were plated in six-well plates. After 24 h, cells were treated with the specified agents. Solvent-treated samples were incubated with 1% (w/v) dimethyl sulfoxide. Cells were collected 72 h later, stained with 0.4% (w/v) Trypan blue solution (1:1, v/v), and counted using a hemocytometer (26).

### 2.4 Ascites collection

Ascites fluids were obtained from women with ovarian cancer or liver cirrhosis undergoing peritoneocentesis for clinical reasons in Ziv Medical Center (Safed, Israel) Helsinki ethics committee approval #0044-19-ZIV and Bnai Zion Medical Center (Haifa, Israel) Helsinki ethics committee approval #0112-20-BNZ and bought from MIDGAM, Israeli biorepository network for research. Patients’ details are summarized in Table 1. All ascites were centrifuged at 3,000 rpm, for 5 min to remove cells. Fluid was aliquoted and frozen at -80°C for future use.

**Table 1.** Ovarian Cancer patients demographic and clinical characteristics.

| Code* | Source | #Case | Age (years) | Histology | Primary/reoccurrence | Treatment | Chemo-resistance |
| --- | --- | --- | --- | --- | --- | --- | --- |
| As001 | Ziv | 10705302 | 79 | tubulovillous adenoma with low grade epithelial dysplasia | primary | before | Yes |
| As002 | Ziv | 10736610 | 68 | N/A | reoccurrence | after (N/A) | Yes |
| As003 | Ziv | 10754148 | 65 | N/A | reoccurrence | after (N/A) | Yes |
| As004 | Midgam | 106936 af-97 | 60 | ovarian carcinoma | N/A | N/A | Yes |
| As005 | Midgam | 106169 af-89.a | 47 | ovarian carcinoma | N/A | N/A | Yes |
| As006 | Midgam | 106866 af-96 | 65 | Adenocarcinoma | N/A | N/A | Yes |
| As007 | Midgam | 107485 af-100 | 66 | high grade serous carcinoma, component of clear cell carcinoma. | N/A | after (N/A) | Yes |
| As008 | Midgam | 107332 af-70a | 66 | Adenocarcinoma | N/A | before | Yes |
| As009 | Ziv | 10831229 | 74 | N/A | reoccurrence | after (N/A) | Yes |
| As010 | Ziv | 10840339 | 75 | N/A | reoccurrence of As009 | after (N/A) | Yes |
| As011 | Bnai Zion | 10686665 | 68 | serous papillary carcinoma | primary | before | No |
| As012 | Bnai Zion | 10690287 | 73 | serous papillary carcinoma | primary | before | Yes |
| As013 | Bnai Zion | 10695219 | 71 | serous papillary carcinoma | primary | before | No |
| Cr001 | Ziv | 10739714 | 79 | N/R | N/R | N/R | Yes |
| Cr002 | Ziv | 10746330 | 82 | N/R | N/R | N/R | No |
| Cr003 | Ziv | 10779177 | 78 | N/R | N/R | N/R | No |
| Cr004 | Ziv | 10785352 | 59 | N/R | N/R | N/R | No |

Abbreviations: \*As denotes OC ascites, Cr denotes cirrhosis ascites; N/A, not applicable; NR not relevant

### 2.5 Ascites experiments

Different concentrations of malignant ascites fluids were applied onto cisplatin resistant OC cell line A2780cisR prior to 5 μM RJY13 exposure. Human A2780cisR cells were plated at 25 × 10^3^ cells/cm^2^ in 25 cm^2^ cell-culture flasks and incubated in 2.5 mL culture medium for 24 hours. 50% conditioned medium (CM) taken from tumor activated HS-5, was used as positive control for chemoresistance induction. Diluted ascites (to final flask concentration of 20% and 30%) were added. RJY13 induced resistance was determined using cPARP immunoblotting Different drugs were applied 5 μL of each (diluted in DMSO), supplemented with the ascites fluid to final concentrations of: 6 μM fisetin, 10 μM 2HE2, 2 μM ruxolitinib, 2 μM tofacitinib, 5 μM dasatinib and 2 μM crizotinib. 5 μL of DMSO vehicle control (0.1%) was added. After 24 hours’ cells were treated with 5 μM RJY13 (final flask concentration) and incubated for additional 24 hours. Cells were collected by using trypsin in RPMI medium, then centrifuged at 3,000 rpm (1,000× g) for 5 min and washed twice with cold phosphate buffered saline (PBS) to obtain cell pellets. Pellets were used to prepare lysates for immuno-blotting or monitoring cPARP by ELISA assay.

### 2.6 Immunoblotting

Immunoblotting was performed as previously described (27, 28). Briefly, Western blot protocol was used for protein analysis on an 8–12% acrylamide gel. Cell lysate samples were prepared for loading by adding lysis buffer (#9803 Cell Signaling Technology, MA, USA) containing protease inhibitors (P8340 and P5726, Sigma, Germany) and phosphatase inhibitor (P-1517, AG Scientific, CA, USA) to the cell pellets. After 30 min, samples were centrifuged and supernatants were tested for protein concentration using DC™ Protein Assay (Bio Rad, USA) by determining absorbance at 630 nm. Samples were lysed in lysis buffer and 50–60 μg protein per monoculture sample, or 100–120 μg protein per co-culture sample were loaded on the gel. Proteins were immunoblotted onto a nitrocellulose membrane (Schleicher & Schuell BioScience GmbH, Germany) which was then blocked with 5% skim milk TBS/T, and incubated with the following antibodies: anti-cleaved poly-ADP-ribose polymerase (PARP) (Asp214) (D64E10, Cell Signaling Technology), anti-α-tubulin (sc-8035, Santa Cruz, TX, USA), anti-GAPDH (#2118, Cell Signaling Technology) and anti-phospho-ERK1/2 (Thr202/Tyr204) (D13.14.4E, Cell Signaling Technology) according to the manufacturers’ instructions. Secondary antibodies,

HRP-linked anti-rabbit (#7074, Cell Signaling Technology) and anti-mouse (NB7539 Novus, Centennial, CO, USA) were used according to the manufacturers’ instructions. Chemiluminescence was performed with SuperSignal™ West Pico PLUS Chemiluminescent Substrate (Thermo Fisher Scientific, MA, USA) and imaged using an HP imager. Densitometry was performed with ImageQuant v8.2 software.

### 2.7 Magpix magnetic beads multiplex

CM samples were diluted 1/2, 1/5, 1/10 in universal assay buffer (in duplicates and tested for IL-1α, IL-1β, IL-5, IL-6, IL-8, IL-10, IL-13, IL-22, G-CSF, INF-γ, TNFα, RANTES (CCL5), leptin and MCP-1 (CCL2) using 96-well ProcartaPlex human multiplex (Thermos Fisher, USA) according to manufacturer instructions. The 96-well plate was analyzed by MAGPIX (Luminex, USA) following the instrument operation procedure.

### 2.8 PathScan® Cleaved PARP Assay

Cleaved PARP as an apoptotic marker in cell lysates was measured using PathScan® Cleaved PARP (Asp214) Sandwich ELISA Kit (Cell Signaling Technology) according to the manufacturer instructions. This ELISA kit is able to detect cPARP from human and murine origin.

### 2.9 Cytokine Array

The human XL cytokine array kit (R&D Systems, Minneapolis, Minnesota, USA) was used to detect relative levels of 105 different human cytokines, chemokines, growth factors and adipokines. This immunoassay utilizes captured antibodies spotted (in duplicate) on a nitrocellulose membrane. Specific proteins bind to antibodies and are detected via biotinylated detection antibodies (sandwich) and visualized using chemiluminescence reagents. The signal produced is proportional to the amount of analyte bound.

### 2.10 Metabolites determination- Sample preparation

The different ascites samples were aliquoted into 300 µL triplicate and frozen at -20°C. In preparation for LC-MS/MS samples were thawed on ice and extracted by dissolving with 1200 µL methanol LC-MS grade. Following 2 min vortex, samples were centrifuged at 13,000 rpm, 4°C, for 15 min. Extracts were transferred into new tubes and evaporated with nitrogen, then dissolved with 150 µL of 50% methanol in double distilled water (DDW) and centrifuged at 13,000 rpm, 4°C, for 10 min. 120 µL of the extracts were transferred into 1.5 ml vials with insert suitable for HPLC and LC-MS/MS. Quality control (QC) samples were prepared by mixing 20 µL from each extract. Methanol was used as blank.

### 2.11 HPLC analysis

The samples were analyzed by injecting 5 μL of the extracted solutions into a UHPLC connected to a photodiode array detector (Dionex Ultimate3000), with a reverse-phase column (ZORBAX Eclipse Plus C18, 100*3.0 mm, 1.8μm). The mobile phase consisted of (A) DDW with 0.1% formic acid and (B) acetonitrile containing 0.1% formic acid. The gradient was started with 98% A and increased to 30% B in 4 min, then increased to 40% B in 1 min and kept isocratic at 40% B for another 3 min. The gradient increased to 50% in 6 min, and increased to 55% in 4 min and finally increased to 95% in 5 min and kept isocratic for 7 min. phase A returned to 98% A in 3 min and the column was allowed to equilibrate at 98% A for 3 min before the next injection. The flow rate was 0.4 mL/min.

### 2.12 LC–MS/MS analysis

MS/MS analysis was performed with Heated Electrospray ionization (HESI-II) source connected to a Q Exactive™ Plus Hybrid Quadrupole-Orbitrap™ Mass Spectrometer Thermo Scientific ™. ESI capillary voltage was set to 3000 V, capillary temperature to 350°C, gas temperature to 350°C and gas flow to 35 mL/min. The mass spectra (m/z 100–1000) acquired in Negative and Positive-ion mode with high resolution (FWHM=70,000). For MS 2 analysis collision energy was set to 30, 50 and 70 EV.

### 2.13 Data preprocessing

Peak determination and peak area integration performed with Compound Discoverer 3.3 (Thermo Xcalibur, Version 3.3.0.305). Auto-integration was manually inspected and corrected if necessary. For some of the compounds, identification was based on MzCloud database using MS 2 data and ChemSpider database using MS 2 data and HRMS.

### 2.14 Statistical analysis

Peak area medians, standard deviation (SD) and relative SD (RSD) were calculated by the Compound Discoverer 3.3 program (Thermo Xcalibur, Version 3.3.0.305). Significant differences between groups were calculated using one-way ANOVA model with Tukey as post-hoc test. Peak areas were normalized using pool samples injected.

## 3 Results

### 3.1 Malignant OC ascites fluids promote platinum chemoresistance of OC cell lines

Ascites fluid has been linked clinically to poor prognosis and chemoresistance (3), therefore the present study monitored the ability of ascites fluids to promote platinum chemoresistance to OC cells. For this research, ascites derived from OC patients was separated into cells and liquid fluid. Non-malignant ascites derived from cirrhotic patients served as a control. The fluid was tested for its ability to promote platinum chemoresistance to OC cell lines by exposing OC cells to different amounts of ascites fluids in the presence of a platinum compound. OC cells were subjected to ascites fluids at concentrations of 10%, 20%, 25%, 30%, and 40% after which their sensitivity to platinum (RJY13) compounds was examined (Fig. 1). In this study, we used the cisplatin-resistant and RJY-13-sensitive OC cell line A2780cisR (25).

**Figure 1.**
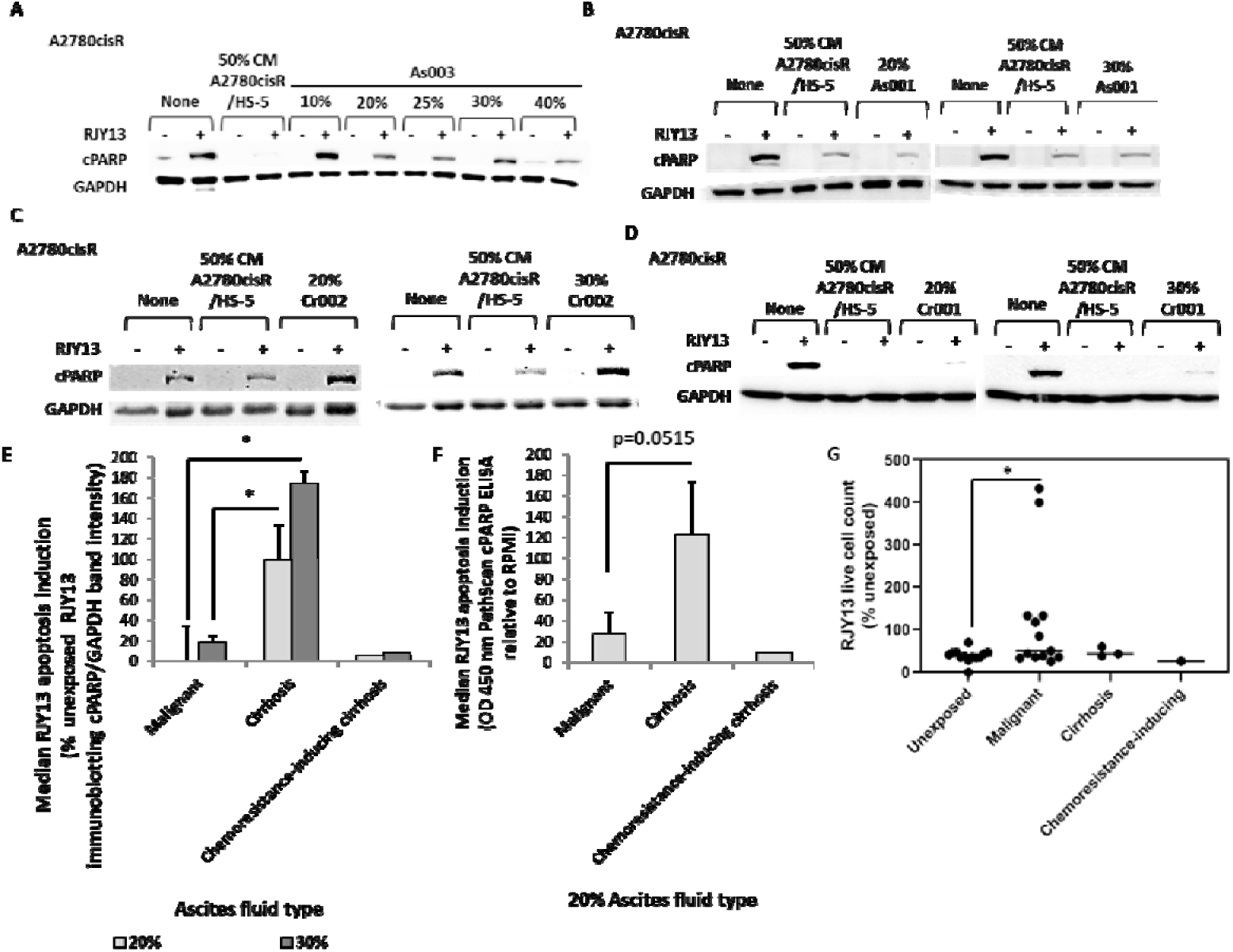
Malignant ascites fluid promote RJY13 chemoresistance in A2780cisR cell lines. 50% conditioned medium (CM) taken from tumor activated HS-5, was used as positive control for chemoresistance induction. RJY13 induced resistance was determined using cPARP immunoblotting (A). 20% (and in some cases also 30%) of ascites fluids were applied to test the ability to trigger chemoresistance in A2780cisR: malignant ascites fluid (B), cirrhosis ascites fluid (C) and the one cirrhosis fluid that conferred chemoresistance to A2780cisR cell line (D). To determine apoptosis and chemoresistance, levels of cPARP were evaluated by immunoblotting and densitometry was quantified. Median cPARP/GAPDH band density relative to control RJY13-treated was calculated (E); malignant ascites n=13, cirrhosis ascites n=3, chemoresistance-inducing cirrhosis ascites n=1, GAPDH was used as loading control. * p<0.05. cPARP was also detected using PathScan cPARP ELISA kit (Cell Signaling Technology, Danvers, Massachusetts, USA) and median OD 450 nm relative to control platinum-treated cells was calculated (F); malignant ascites n=6, cirrhosis ascites n=3, chemoresistance-inducing cirrhosis ascites n=1. Bars indicate standard deviation. (G); resistance was also estimated by live cell counts relative to cells unexposed to ascites and untreated; malignant ascites n=13, cirrhosis ascites n=3, chemoresistance-inducing cirrhosis ascites n=1.

Results shown in Figure 1A indicate that maximal chemoresistance was induced in A2780cisR cells by the addition of 20-25% of malignant ascites fluid, therefore 20% (and 30% for part of the fluids) was used to investigate their ability to monitor chemoresistance using both immunoblotting and cPARP ELISA assays. Figure 1B shows an example of the chemoresistance promoted in A2780cisR cells by 20% and 30% of malignant ascites As001.

A total of 11 malignant ascites out of 13 samples tested, derived from different ovarian cancer patients (Table 1), were active in conferring chemoresistance with variable potency as monitored by the reduction of cPARP levels in A2780cisR cells treated with platinum IV prodrug (Fig. 1B and 1E). In contrast, most of the cirrhosis ascites (3 out of 4 ascites) failed to promote platinum chemoresistance to OC cells (Fig. 1C, 1E). Interestingly, one cirrhosis ascites (Cr001) was active in promoting platinum chemoresistance to OC cells (Fig. 1D).

Ascites fluid derived from OC patients promoted chemoresistance causing moderate (20%) and significant (30%) reduction in cPARP levels as summarized in Figure 1E. The above-mentioned results were confirmed by utilizing cPARP PathScan ELISA to monitor levels of cPARP (Fig. 1F). Low median cPARP signal obtained, when OC cells were exposed to malignant ascites and chemoresistance-inducing cirrhosis ascites fluids, indicated that A2780cisR cells exhibit low sensitivity to RJY13. On the other hand, high cPARP signal was observed in A2780cisR cells pre-treated with fluid derived from cirrhosis patient, indicating no reduction in RJY13 sensitivity. Exposure to malignant ascites resulted in the elevation of cell counts in A2780cisR cells treated with RJY13 compound, while ascites derived from cirrhosis patients did not result in an increase in cell count (Fig. 1G). Interestingly, the one cirrhosis ascites fluid (Cr001) that mediated RJY13 resistance to A2780cisR cells, resulted in elevation of cPARP levels.

### 3.2 Elevated levels of IL-6 in malignant ascites are a primary suspect responsible in mediating platinum chemoresistance in A2780cisR cell line

The content of soluble factors in the different ascites were analyzed by the cytokine array (R&D Systems, Minneapolis, Minnesota, USA). Results in Figure 2 showed the dot density (Fig. 2A) normalized to reference dots (Fig. 2B). The dot density ratio of the malignant to cirrhosis ascites was calculated (Fig. 2C) showing soluble factors enriched in the ascites fluids that were active in conferring platinum chemoresistance. Manhattan distance for that ratio with the relevant *p*-value for each of the soluble factors examined was computed (Fig. 2B). In addition, multiplex ELISA (MagPix) was utilized to monitor levels of IL-1α, IL-1β, IL-5, IL-6, IL-8, IL-10, IL-13, IL-22, G-CSF, INF-γ, TNFα, RANTES (CCL5), leptin and MCP-1 (CCL2) concentrations (Luminex Corporation, Austin, Texas, USA).

**Figure 2.**
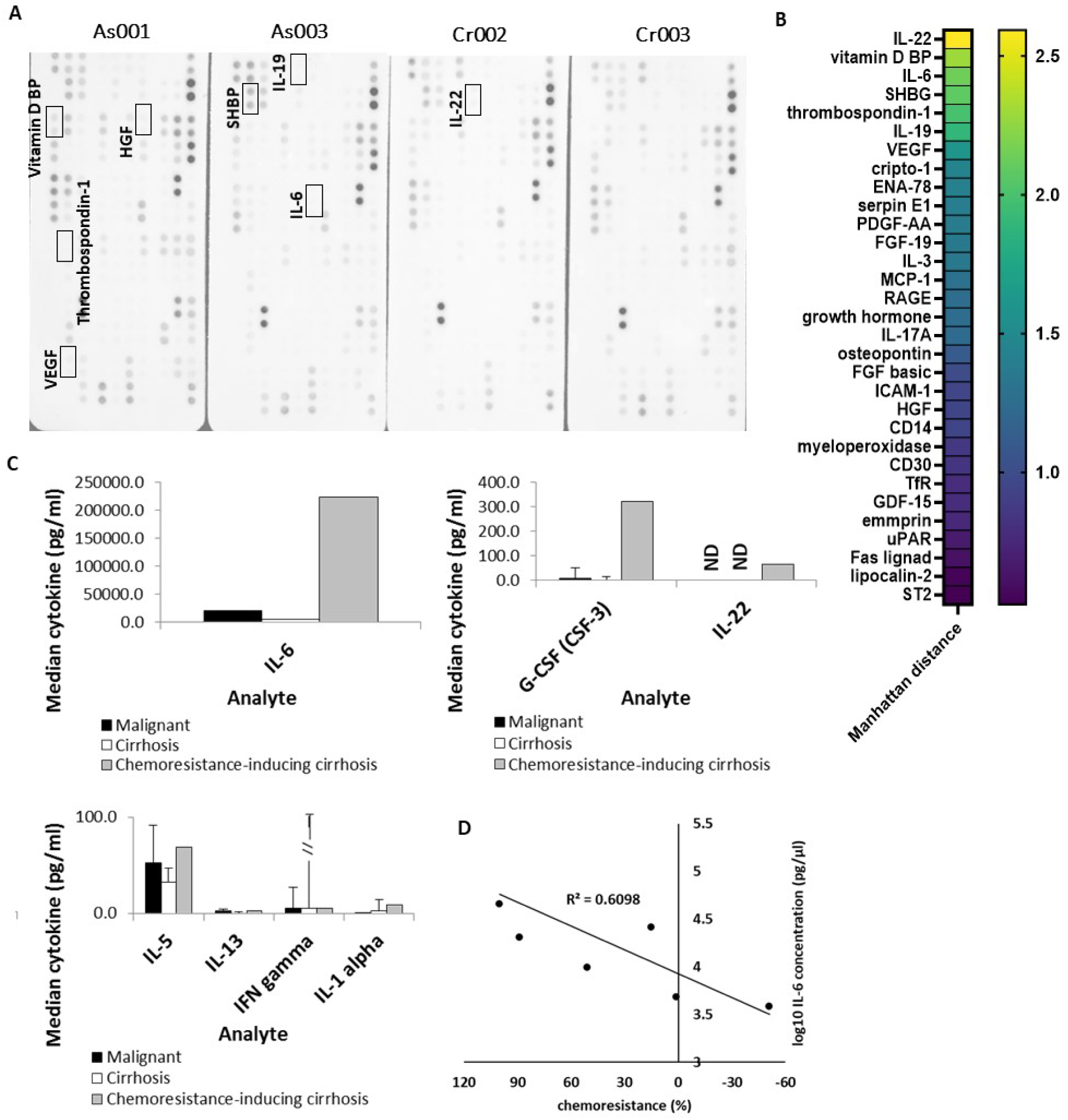
Identification of soluble factors enriched in malignant and cirrhosis ascites. Ascites fluid derived from OC and cirrhosis female patients was tested to detect levels of different soluble factor using human XL cytokine array kit (R&D Systems, Minneapolis, Minnesota, USA). (A) Dot density was measured and malignant ascites to cirrhosis ascites ratio was calculated for detectable factors. Manhattan distance was calculated for this ratio (B) to present significant differences between malignant ascites (n=2) and cirrhosis ascites (n=2). Multiplex ELISA kit (Thermo Fisher Scientific, Paisley, UK) designed to detect presence of enriched cytokines (IL-1α, IL-1β, IL-5, IL-6, IL-8, IL-10, IL-13, IL-22, G-CSF, INF-γ, TNFα, RANTES (CCL5), leptin and MCP-1 (CCL2) was used in 3 ascites groups, according to their ability to induce chemoresistance in A2780cisR cells: malignant ascites (n=3), cirrhosis ascites (n=3) and chemoresistance-inducing cirrhosis ascites (n=1) (C). IL-6 concentration in each of the ascites fluid tested was compared to its relative ability to confer RJY13 platinum resistance (D).

In Figure 2B, IL-22 is shown as the most prominent cytokine, found in higher concentrations in the cirrhosis ascites. In addition, Vitamin D BP, and to a lesser extent IL-6, IL-19, VEGF and G-CSF (Fig. 2C) were found as prominent factors in the OC ascites fluid (Fig. 2B and 2C). IL-3, MCP-1, growth hormone, IL-17A, basic FGF, HGF and others were also enriched in the OC ascites (Fig. 2B).

Interestingly, the cirrhosis ascites that was active in promoting chemoresistance was also enriched with IL-6 and G-CSF (Fig. 2C). However, IL-6 levels in malignant ascites were higher than the levels found in cirrhosis ascites fluid, though still significantly lower than the cirrhosis ascites fluid that was active in promoting chemoresistance to OC cells.

Figure 2D compares IL-6 concentration (logarithmic scale) to RJY13 sensitivity (calculated as 100%-(apoptosis rate). RJY13 sensitivity was measured by cPARP immunoblotting, for each of the ascites fluid individually, except for the ascites fluid derived from one of the cirrhosis patients (Cr001). Figure 2D exhibited a good correlation between the levels of IL-6 as determined in the various ascites fluids and the observed sensitivity to RJY13. In general, high levels of IL-6 correlate with low sensitivity to RJY13. Thus, the levels of IL-6 seem to serve as an indicator of RJY13 resistance in OC cells.

### 3.3 JAK/STAT inhibitors increase RJY13 sensitivity in resistant OC cells pre-treated with malignant ascites fluid

IL-6, the cytokine found to correlate with induction of chemoresistance in A2780cisR by ascites fluid is known to activate the JAK/STAT signal transduction pathway (29). Two FDA-approved JAK inhibitors: Ruxolitinib (JAK1/2 inhibitor) (30) and Tofacitinib (JAK1/3 inhibitor) (31) were used. Dasatinib (Src and Abl inhibitor) was used as well (32). These kinase inhibitors were applied together with the ascites fluid, 24 hours prior to RJY13 treatment. Results presented in Figure 3 showed that OC cells that pre-treated with malignant ascites fluids and exposed to combination of JAK inhibitors such Ruxolitinib, Tofacitinib or Dasatinib with platinum compounds, resulted in partial restoration of platinum sensitivity as demonstrated by the elevated levels of cPARP band density in treated OC cells (Fig. 3A).

**Figure 3.**
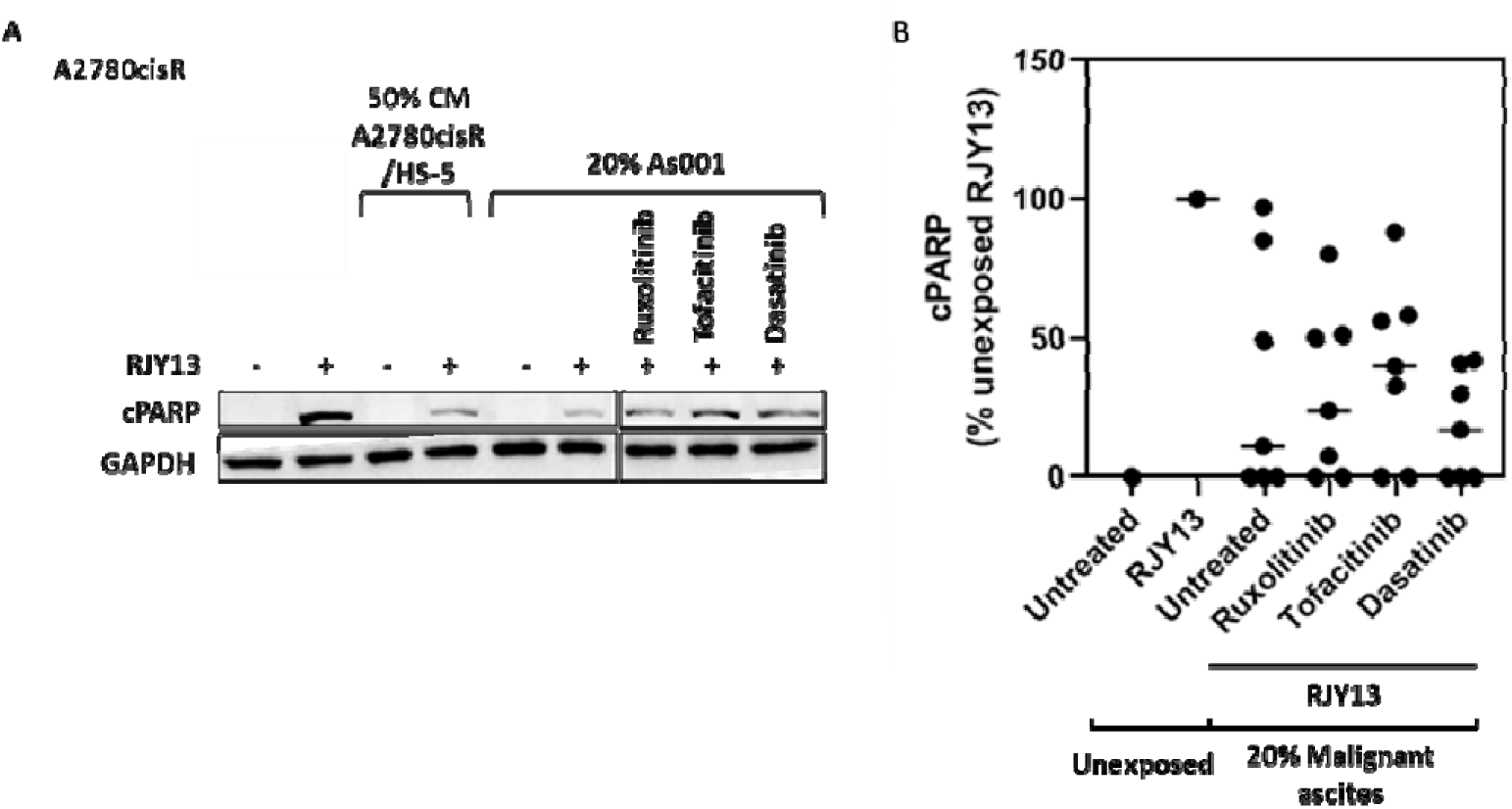
JAK inhibitors partially restore chemo sensitivity to OC cells mediated by malignant ascites-fluids. Malignant ascites fluids (OC patients) and non-malignant ascites fluids (Cirrhosis patients) were incubated with cisplatin-resistant OC cell line A2780cisR together with one JAK inhibitor (2 μM Ruxolitinib, or 2 μM Tofacitinib or 5 μM the multi-kinase inhibitor Dasatinib), combined with 5 μM RJY13 for 24 hrs. 50% CM of tumor activated HS-5 were used as positive control (A). The levels of cPARP were evaluated by immunoblotting and quantified by densitometry. Median cPARP/GAPDH band density relative to control for the kinase inhibitors are indicated (B). Malignant ascites untreated (n=8), cirrhosis ascites (n=3) and chemoresistance-inducing cirrhosis ascites (n=1). GAPDH was used as a loading control.

The OC cells were exposed to the ascites fluid from the different groups; malignant, cirrhosis and cirrhosis fluid that induced chemoresistance (Cr001). We monitored median relative apoptosis (cPARP/GAPDH immunoblotting) levels in the presence and absence of JAK inhibitors (Ruxolitinib or Tofacitinib) and Dasatinib. A moderate effect was observed when the JAK inhibitor was combined with the platinum compound, but less when Dasatinib was used. Interestingly, Tofacitinib was more potent than Ruxolitinib in restoring platinum sensitivity to OC cells (Fig. 3B).

### 3.4 Crizotinib promotes RJY13 sensitivity in resistant OC cells pre-treated with malignant ascites fluid

Although IL-6 was shown to be enriched in malignant ascites, inhibition of IL-6 function resulted in moderate and partial restoration of platinum sensitivity, which argues for the possible involvement of additional factors (such as HGF or others) in mediating chemoresistance.

Previously we showed that Crizotinib (HGF/c-Met inhibitor) was shown to overcome platinum chemoresistance mediated by tumor activated HS-5 (TA HS-5) conditioned medium (CM) (23). The current study shows that HGF was also enriched in malignant ascites and thus its involvement in promoting platinum chemoresistance was examined.

Results shown in Figure 4A demonstrated that the presence of 20% malignant ascites fluid resulted in reduction in cPARP levels (Fig. 4A), implying induction of platinum chemoresistance. The ability of Crizotinib, a multi-kinase inhibitor targeting c-MET, ALK, ROS1 and other kinases (22), to affect platinum chemoresistance in OC cells was investigated. The current results revealed that exposure to Crizotinib caused a significant increase in cPARP levels, implying the restoration of platinum sensitivity. Conversely, Crizotinib was also found as potent in restoring platinum chemoresistance mediated by malignant ascites in 3 malignant ascites tested (Fig. 4B).

**Figure 4.**
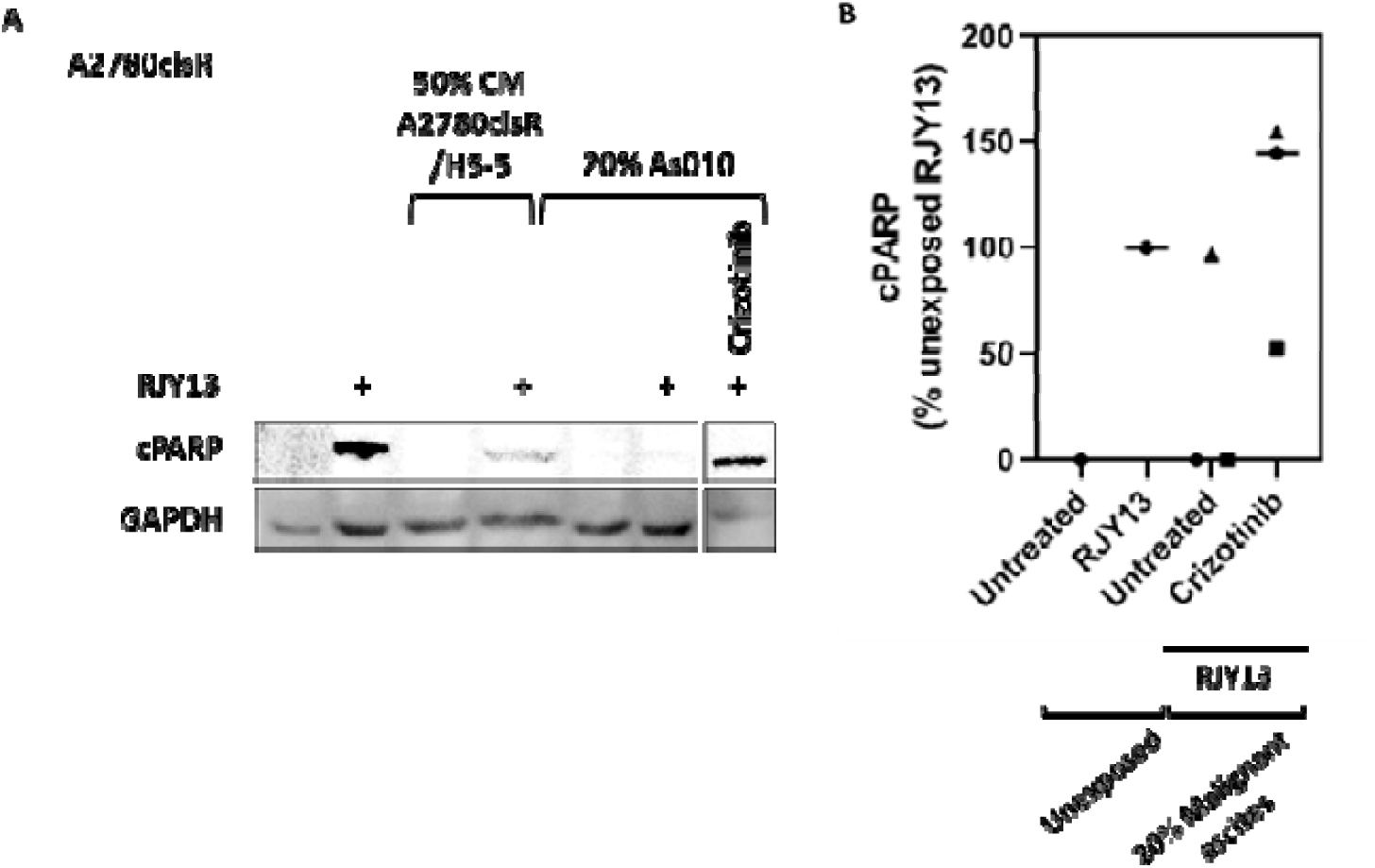
Crizotinib restores RJY13 malignant ascites mediated chemosensitivity to OC cells. Cisplatin-resistant OC cell line A2780cisR was incubated with 20% ascites fluid derived from OC patients with or without Crizotinib (2 μM) for 24 hours. Exposure to 50% conditioned medium (CM) of tumor activated HS-5 was used as positive control (A). Levels of cPARP were evaluated by immunoblotting and quantified by densitometry. Median cPARP/GAPDH band density relative to control was calculated (B). Untreated (n=3), Crizotinib (n=3). GAPDH was used as a loading control.

### 3.5 2-hydroxy estradiol (2HE2) elevates RJY13 platinum sensitivity in a resistant OC cell line pre-treated with malignant ascites fluid

The estradiol derivative 2HE2 was demonstrated to be a potent agent that restores platinum chemosensitivity to OC cells, mediated by cellular and soluble compartments of TME in *in-vitro* systems (21). Thus, we investigated its ability to restore platinum sensitivity to OC exposed to malignant ascites.

The results in Figure 5A show that a significant decrease in cPARP levels is obtained when A2780cisR cells were exposed either to CM derived from TA HS-5 or from malignant ascites derived from OC patients. Interestingly, addition of 2HE2 significantly restored platinum sensitivity as evident by increasing levels of cPARP (Fig. 5A). cPARP band density was quantified for several OC ascites (n=6) and cPARP/GAPDH -ratio for each sample were shown in the presence or absence of 2HE2 (Fig. 5B). The results show a significant increase in the ratio of cPARP/GAPDH in 2HE2 treated samples, implying a restoration of platinum sensitivity to OC cells. Interestingly, 2HE2 was active in restoring platinum sensitivity in every malignant ascites sample used in this study.

**Figure 5.**
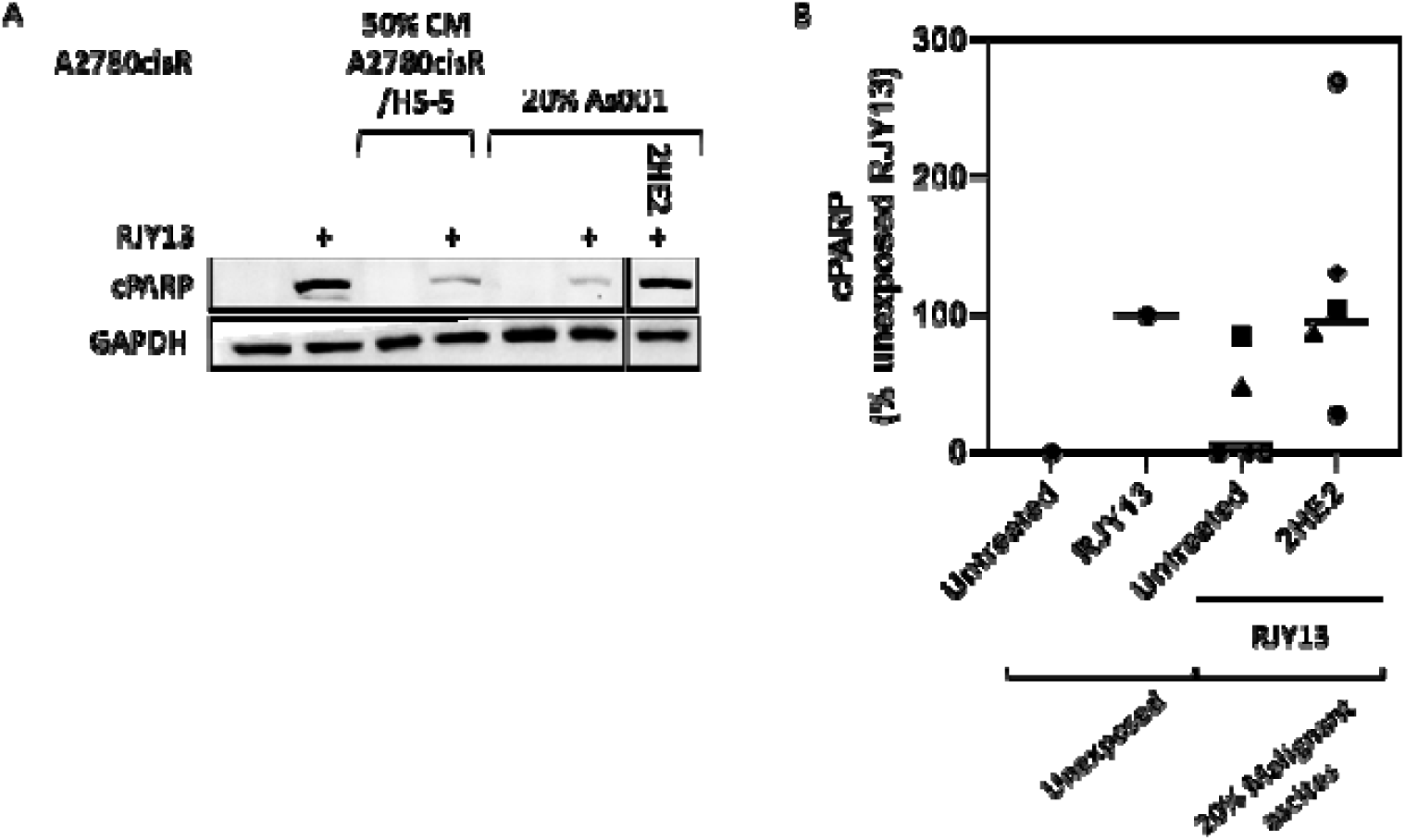
2HE2 restores RJY13 chemosensitivity to OC cells that mediated by malignant ascites soluble factors. A2780cisR, cisplatin resistant OC cells were treated with 5μM RJY13 in the absence of 50% tumor activated HS-5 CM or 20% As001 (A). Also shown sample of A2780cisR cells exposed to 20% of As001 ascites fluids in the combination of 10μM of 2HE2 (A). The levels of cPARP were evaluated by immunoblotting and quantified by densitometry. Median cPARP/GAPDH band density relative to control is indicated (B). Malignant ascites (n=6). GAPDH was used as a loading control.

Unlike malignant ascites, most of cirrhosis ascites were unable to promote platinum chemoresistance to OC cells. However, one cirrhosis ascites (Cr001) was able to promote platinum chemoresistance to OC cells. Interestingly, the Cr001 capable of inducing platinum chemoresistance was enriched in several growth factors such as GCSF and Il-22 (Fig. 3C). Thus, the present study inquired whether the modulators used above have the ability to restore platinum sensitivity to OC cells exposed to Cr001 ascites.

Figure 6 shows, as expected, that exposure of OC cells to Cr001 ascites fluid confer platinum resistance (Fig. 6A). Combination of modulators (Fig. 6A and 6B) with platinum compounds resulted in various responses. Combination of RJY13 with Fisetin, 2HE2, Ruxolitinib, Tofacitinib, and Dasatinib (Fig. 6A) was not active in restoring platinum sensitivity to OC cells exposed to Cr001 ascites. In contrast, combination of RJY13 with Crizotinib resulted in restoring platinum sensitivity to OC cells (Fig. 6B) similar to malignant ascites As001 and As012 (Fig. 6B).

**Figure 6.**
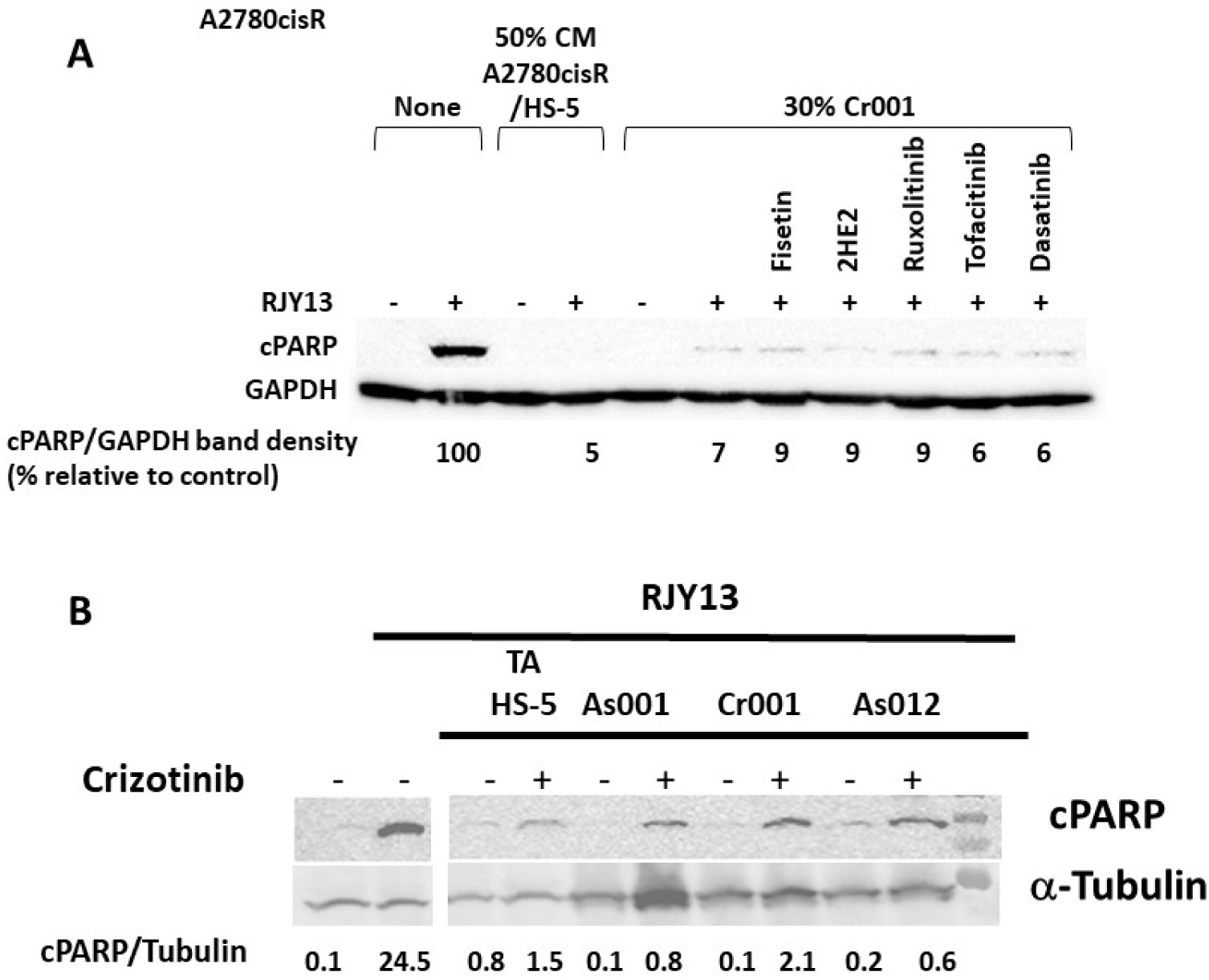
Crizotinib restores platinum sensitivity to OC exposed to cirrhosis ascites (Cr001). A2780cisR, cisplatin resistant OC cells were treated with 5μM RJY13 in the absence of CM, with 50% of tumor activated HS-5 CM and 30% of Cr001 combined with 10μM of Fisetin, 10μM 2HE2, 2μM of each Ruxolitinib, Tofacitinib or Dasatinib (A). A2780cisR cells were treated with 5μM RJY13 in the presence of the appropriate CM derived from TA-HS-5 or ascites fluids (As001, Cr001, and As012). Crizotinib at 2μM was combined in the appropriate samples as indicated (B). Levels of cPARP were evaluated by immunoblotting through quantified densitometry. Median cPARP/housekeeping protein (GAPDH or α-tubulin) band density relative to control is indicated (A, B).

### 3.6 Metabolites that are enriched in malignant ascites

In this study we focused on the role of soluble growth factors in mediating platinum chemoresistance to OC cells. However, the ascites fluids contain other materials besides soluble growth factors, such as different metabolites, exosomes, etc. (33). Thus, metabolites in malignant ascites as well as in cirrhosis ascites were monitored using LC/MS-MS. Compounds were identified using the mZcloud database. The ratio for normalized area of the malignant to the cirrhosis ascites fluids, *p*-values and Manhattan distance (|log2(As/Cr)|+| log10(*p*-value)|) were calculated for each detected compound. The results shown in Figure 7 identified a number of metabolites, that are of special interest due to their involvement in the arachidonic acid (AA) pathway (Fig. 7B)

**Figure 7.**
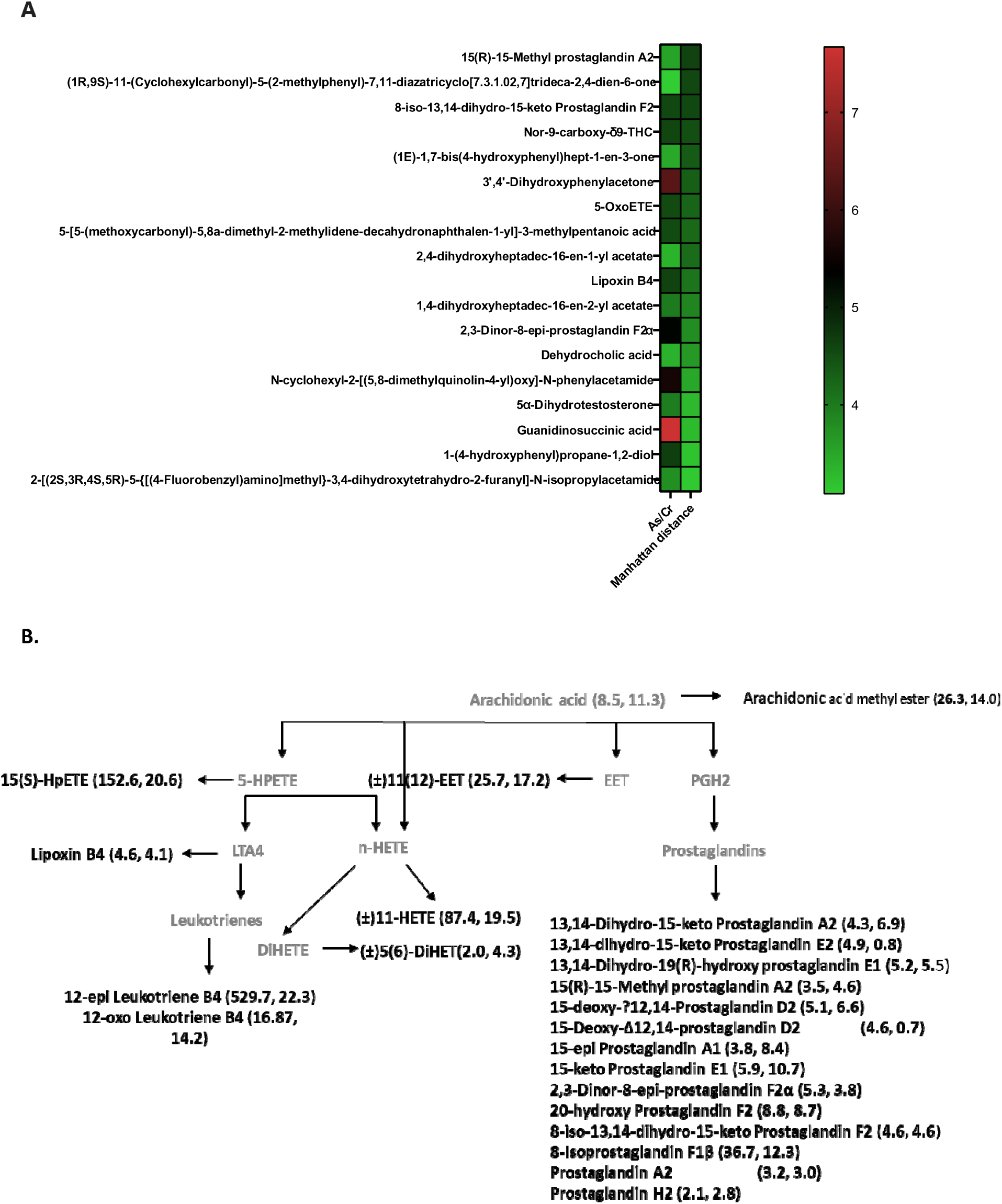
Identification of metabolites that are enriched in malignant ascites and cirrhosis ascites fluid. (A) Ascites fluids from OC and cirrhosis -female patients were analyzed for metabolites using LC-MS/MS. The fluids were divided into 2 groups (1: As001-As003, Cr001 and 2: As004-As013, Cr004) and tested separately. Compounds were identified using mZcloud data base and the median ratio of the normalized area of the malignant ascites to the cirrhosis ascites was calculated for each compound. *p*-value of the malignant vs the cirrhosis ascites was determined (fold change >3) and Manhattan distance was calculated. Threshold Manhattan distance>3 (and *p*-value<0.01) was set. Drugs, food supplements and food components, cosmetic and cleaning agents, herbicides and plastic monomers contaminants were removed from the list. Malignant ascites (n=13), cirrhosis ascites (n=2). (B) Metabolites of arachidonic acid showing fold changes between malignant ascites (As) vs. cirrhosis ascites (Cr). Malignant ascites (n=9, As004-013), cirrhosis ascites (n=1, Cr004).

Fatty acids and fatty acids derivatives were found in both types of ascites; however, they were enriched in the malignant ascites. Detailed analysis is shown in the supplemental materials (Figure S1). Figure 7 shows abundance of metabolites of the AA pathways: AA and its metabolites such as arachidonic acid methyl ester, as well as cytochrome P450 (CYP450), lipooxegenases, ω-hydroxylases, epoxygenases and cyclooxygenases (COXs) products: hydroxyeicosatetraenoic (HETE), hydroperoxyeicosatetraenoic (HPETE), epoxyeicosatrienoic (EET) acids and prostaglandins (PG).

Another fundamental group of substances (which is also related to the AA group) observed in the malignant ascites consists of DHA, EPA and their CYP450, lipoxygenase and COX metabolites (data not shown). In addition, several amino acids metabolites were also found in higher concentration in the malignant ascites, their significance and their role in promoting drug resistance and malignancy remains to be investigated.

## 4 Discussion

This study is a follow-up to our earlier research that showed how a TME component promotes OC chemoresistance (23). Here, we carried out ex vivo assays to track the capacity of malignant ascites derived from OC patients to promote chemoresistance in OC cells. We compared malignant ascites to ascites derived from cirrhosis patients. Due to the small number of ascites samples examined—13 malignant ascites and 4 cirrhosis ascites—we view this study as a pilot study. However, we believe that the results of our study offer insightful information and helps to clarify how malignant ascites affects platinum chemoresistance. Our findings, as shown in Figure 1, demonstrated that the preponderance of malignant ascites— more than 92%—were active in conferring platinum chemoresistance to OC cells. In contrast, only one of the four cirrhosis ascites was active in conferring OC platinum chemoresistance, while the other three were inactive (Fig. 1).

In this research, the soluble component of ascites fluids was used. Malignant ascites has the ability to promote chemoresistance, in contrast to most cirrhosis ascites, which lead to identify soluble growth factors that are enriched in the active ascites promoting chemoresistance as compared to non-active ascites.

The ability of MSC to promote platinum chemoresistance to OC in a manner dependent on extracellular signal-regulated kinase (ERK1/2) has been reported previously (21). The involvement of ERK1/2 in soluble factor-mediated chemoresistance was inconclusive and was a cell type-specific phenomenon (23).The current study investigated the possible role of ERK1/2 activity in platinum chemoresistance mediated by OC. There was no correlation between the status of ERK1/2 phosphorylation and the presence of different types of ascites (data not shown).

The amounts of the various growth factors in 2 malignant ascites that were active in promoting OC chemoresistance were compared to 2 cirrhosis ascites that were not active in promoting OC chemoresistance by using cytokine arrays (Fig. 2). The findings revealed that the cancerous ascites had higher concentrations of a number of growth factors, including IL-6, IL-19, G-CSF, HGF, and others.

Intriguingly, IL-22, a cytokine produced by various immune cell populations at an inflammatory location, was enriched in cirrhosis ascites. Elevated IL-22 has been linked to decreased survival from complications related to the prognosis of advanced liver cirrhosis (34).

One of the enriched growth factors in malignant ascites that might be responsible for causing platinum chemoresistance was the cytokine IL-6 (Fig. 2). The present study findings, is supported by previous report which linked the presence of IL-6 in ascites fluids to platinum resistance (35). Both tumor cells and TME cells secrete IL-6 (11, 36). Paclitaxel drug resistance was induced in primary OC cells by cancer associated fibroblasts (CAFs) secreting IL-6 via the JAK2/STAT3 signal transduction pathway (11). In breast cancer, MSC secreted IL-6 and CXCL-7 that controlled cancer stem cells (CSC) and accelerated tumor growth (37). Cisplatin resistance was linked to elevated amounts of IL-6 in ovarian cancer cells, both exogenously and endogenously (38). JAK inhibitors that prevent IL-6 activity were used to examine the significance of elevated IL-6 levels on the ability of malignant ascites in promoting OC chemoresistance (35). The JAK1/2 and JAK3 inhibitors were Ruxolitinib and Tofacitinib, respectively (35,36). Both of these inhibitors only slightly restored platinum sensitivity (Fig. 3). The IL-6/JAK/STAT3 axis is well known to promote chemoresistance in a number of cancer types (39–41). Ruxolitinib, according to pre-clinical research, limits tumor growth in immune-competent mice and sensitizes OC cells to Taxol (42). Comparison of the drugs activity, Ruxolitinib vs. Tofacitinib in restoring platinum sensitivity to OC cells, showed that Tofacitinib was more effective in restoring platinum sensitivity (Fig.3). Although Tofacitinib was not studied in ovarian cancer, its interaction with PI3K may account for its superior effectiveness over Roxolitinib in re-sensitizing OC cells (43). Additionally, based on our preliminary findings, it is advised to monitor Tofacitinib’s possible effectiveness in clinical settings in OC patients.

Platinum sensitivity was not restored by Dasatinib, a multi-kinase inhibitor that targets Abl and Src (Fig. 3). Consistent with clinical observations, the effect of Dasatinib’s as a single agent has a limited effect in patients with recurrent OC (44). Other Src inhibitors, such as Saracatinib, showed minimal toxicity when combined with carboplatin or paclitaxel in a Phase I clinical trial research (45). Intriguingly, we showed that Dasatinib was effective in reversing platinum sensitivity to OC acquired drug resistance as a consequence of direct co-culture with MSC cells (24), but not when chemoresistance was mediated by soluble factors (23). It could be assumed that Dasatinib targeted kinases are not involved in soluble factors mediated chemoresistance in OC cells.

A high level of Hepatocyte growth factor (HGF) was detected in the malignant ascites, despite not being one of the most significant components in the ascites fluids that induce chemoresistance. HGF was associated with advanced stages of OC cancer (46), histology types and poor prognosis in OC (46, 47). Additionally, it was discovered that HGF, which is secreted by OC cells, affects ascites development and renders mesothelial cells to become more conducive to cancer cell invasion (47).

Crizotinib is an ALK, ROS1, and c-MET inhibitor that has been approved by the FDA in treatment of several human cancers (48). According to our previous finding, Crizotinib also inhibits JAK2 (22). In the presence of malignant ascites, Crizotinib was effective in restoring platinum sensitivity to OC (Fig. 4). Crizotinib’s capacity to inhibit multiple kinases, including JAK2 and c-MET, which are implicated in mediating platinum chemosensitivity, may explain its potency in restoring chemoresistance to platinum-based drugs. Overexpression of c-MET was reported in 7–27% of OC patients. Its activation was linked to a poor prognosis in individuals with high-grade serous ovarian cancer (HGSOC) (49), lung, breast, stomach, and head and neck cancers. Treatment with Crizotinib increased apoptosis and reduced cell proliferation in OC cell carcinomas with elevated levels of c-MET. A clinical trial with Crizotinib, treating OC patients with amplified Crizotinib target genes (ALK, MET and ROS proto-oncogene 1 (ROS1)) was initiated (50).

Crizotinib synergized with cisplatin to inhibit ovarian cancer cells growth in vitro and in vivo (51). This suggest that Crizotinib might potentiate the activity of cisplatin in ovarian cancer. Moreover, Crizotinib synergizes with PARP inhibitors and carboplatin for enhancement of apoptosis in ovarian cancer (52). Consistent with preclinical research, in a case report study, Crizotinib exhibited effective activity in ovarian cancer patients with ROS1 rearrangement (53).

Similarly, 2HE2, an estradiol derivative devoted of estrogenic activity, was successful in restoring platinum sensitivity to OC (Fig. 5). The 2ME2 and its metabolite, 2HE2, have undergone intensive research and have been shown to have strong cardiovascular and anti-cancer effects. While 2ME2’s central challenge is its poor bioavailability, clinical studies utilizing 2ME2 have been conducted not only in ovarian cancer but also in breast and prostate cancer (54). Additionally, 2ME2 was linked to induced cell apoptosis in preclinical experiments by reducing the expression of anti-apoptotic proteins like Bcl-xl, Bcl-2, and c-myc (55). Moreover, was noted that Dasatinib’s ability to induce apoptosis was augmented by 2ME2 (55).

The cirrhosis ascites Cr001 which was active in promoting platinum chemoresistance to OC cells (Fig. 1) was enriched with a number of cytokines and growth factors including IL-6 (Fig. 2 and Fig. 3). Restoring platinum activity to OC cells exposed to Cr001 ascites was possible by combining 2HE2 and Crizotinib (Fig. 6), similar to their ability in restoring platinum sensitivity to OC cells exposed to malignant ascites (Fig. 3-5). The ability of cirrhotic ascites Cr001 to confer platinum chemoresistance to OC cells could be mediated by various growth factors that were enriched in Cr001, such as IL -6, GCSF, and IL -22 (Fig. 3).

Metabolites present in malignant ascites may play a role in OC chemoresistance in addition to growth factors’ role in mediating this phenomenon. To identify metabolites that are prominent in malignant ascites, we used LC-MS/MS (Fig. 7, Fig. S1 and Table S1). A significant number of metabolites (Table S1) were found to be enriched in malignant ascites in comparison to cirrhosis ascites. The significance and the role of theses metabolites remains to be investigated.

Predicting patient response to therapy and the onset of chemoresistance is one of the long-term challenges in cancer biology. The complex tumor microenvironment which includes the metabolic compartment, has a substantial impact on chemoresistance (33). The higher presence of fatty acid metabolites in the malignant ascites than in the cirrhosis ascites can indeed be considered to be among the main components of metabolites in the malignant ascites. Fatty acids can provide a valuable nutrient and energy source for tumor growth. Arachidonic acid (AA), one of the metabolites of fatty acids, is crucial in promoting OC chemoresistance (56). It is interesting to note that poor prognosis and overexpression of fatty acid synthase (FASN) was significantly correlated with OC patients’ tumor grade, stage and poor prognosis. Consequently, inhibition of FASN in cisplatin resistance OC cells resulted in resistance mitigation (57).

Our results are consistent with earlier research that investigated metabolites in malignant ascites (56,57). Punnonen et al., reported higher and substantial amounts of PGE2, leukotriene B4 and palmitoleic acid in malignant ascites fluids relative to the peritoneal fluids of patients with gynecological benign disease (58). Additionally, Low et al. (2022) found aberrant lipid metabolism and lipolysis-promoted activity in the sera and ascites of OC patients (33).

The abundance of prostaglandin (PG) metabolites in malignant ascites, both in terms of quantity and content, is an indication of an inflammatory environment and increased COX activity. PGE2, which is produced in high concentrations in tumors by an altered AA pathway, can stimulate a series of pro-tumorigenic pathways through the protein kinase A (PKA), PI3K/Akt, and Ras/MEK-ERK pathways (33). The Adenomatous Polyposis Coli (APC)/ catenin/TCF pathway, that regulates cell division, angiogenesis, and apoptosis, is a significant depiction of such a system (59). The RAS-mitogen activated protein kinase (MAPK), which assists in cell survival, can also be induced by PGE2 (60). In fact, a number of studies have indicated a beneficial impact of COX-2 inhibitor combinations in improving cancer therapeutic efficacy (61).

In conclusion, our study showed the significance of malignant ascites in mediating chemoresistance to OC cells. IL-6 and HGF, were particularly abundant in malignant ascites. Modulators of signaling pathways associated with IL-6 and HGF enabled platinum sensitivity restoration. This suggests that a combination of the identified modulators (such as Crizotinib) might be considered and further explored as a new therapeutic strategy in ovarian cancer.

## Author Contributions

YC, HK performed the experiments; AA designed the ex vivo and ascites collection, YC and HK analyzed the data, RF, YS, RK collected ascites liquids and analyzed clinical data, JM and JG designed the research study and wrote the paper. All authors read and approved the manuscript.

## Acknowledgments

This project was supported by a grant from the Israeli Ministry of Health to JM (project 3-0000-15095).

